# Minocycline treatment affects astrocyte – microglia – neuron interaction and functional compensation of motor deficits in rat model of combined fluorocitrate and 6-OHDA lesion and early Parkinson’s disease

**DOI:** 10.1101/2025.06.16.659904

**Authors:** Katarzyna Z. Kuter, Agnieszka M. Chwastek, Martyna Paleczna, Justyna Kadłuczka, Barbara Kosmowska, Innesa Leonovich, Wiktoria Maciasz, Tomasz Lenda, Joanna Kula

## Abstract

Prolonged nervous system inflammation and glia activation are among hallmarks of Parkinson’s disease. There are no therapies slowing pathology. Microglia and astrocytes are considered targets for disease modifying strategies. Nigrostriatal neurodegeneration causes locomotor dysfunction but at early stages can be compensated. Interaction between neurons, microglia and astrocytes could be essential for this functional adaptation and neuronal survival in long-term.

The aim was to check **how microglia activation inhibition affects neuron and astrocyte cell death caused by selective toxins and how it affects locomotion and potential for spontaneous functional compensation of motor deficits** at the early stages of Parkinson’s disease.

In a rat model of fluorocitrate (FC)-induced astrocyte death and microglia activation combined with 6-OHDA selective dopaminergic system neurodegeneration we analyzed anti-inflammatory effect of minocycline on each of the cell type and on functional behavioral output.

In result, reduced microglia activation by minocycline probably prevented part of astrocytes from FC-induced cell death. Microglia inhibition caused non-dopaminergic neurodegeneration in a group treated by both neurotoxins but still enhanced compensatory potential to functionally improve walking deficits caused by dopaminergic lesion.

It seems that activation of microglia by dying astrocytes vs dying neurons induced varied mechanisms. Inhibition of strong microglia activation could be protective for astrocytes but microglia is also important for neuronal adaptation, therefore suppression of its activation perturbs structural rebuilding during progressive neurodegeneration affecting functional outcome.

Understanding the relationship between neuronal death, astrocyte loss of function and microglial response could help to identify new, non-neuronal pharmacological target for healing various neurodegenerative diseases.

**Highlights:**

- Astrocyte death affected neuron function but neurodegeneration did not affect astrocyte survival.
- Microglia was differentially activated by death of astrocytes than by neuron degeneration.
- Minocycline treatment decreased morphological signs of microglia activation, probably protected some astrocytes but negatively affected astrocytes in 6-OHDA lesion group.
- Minocycline treatment despite inducing non-dopaminergic neurodegeneration still enhanced compensatory potential to functionally improve walking after combined 6-OHDA lesion and fluorocitrate-induced astrocyte death.

## 1. Introduction

In the course of Parkinson’s disease (PD) progressing dopaminergic neuronal loss and the presence of cytoplasmic inclusions rich in alpha-synuclein in neuronal and some glial cells are accompanied by reactive changes of astrocytes and microglia. Both types of glial cells participate in inflammation (Ayerra et al., 2024). Both, strong as well as minor but prolonged pro-inflammatory activation state can be deleterious to neuronal cells and other cell types. (Butler et al., 2021; Ugalde-Muñiz et al., 2020). The role of neuroinflammation in the pathogenesis of PD is still undetermined. Glia activation has been considered as a downstream response to neuronal damage and alpha-synuclein aggregation, although loss of protective function or gain of toxic role of astrocytes and microglia was found to be also a relevant mechanism (Tremblay et al., 2019). Many studies focus on either neurons alone or microglia or astrocytes independently, while it is important to look at broader tissue context including various cell types interactions.

Microglia physiological function include formation, maintenance and modification of neuronal circuits through pruning and forming of synapses, and contribution to axonal growth and myelination, removal of debris and apoptotic cells. Microglia activation follows a wide spectrum of phenotypes (Fumagalli et al., 2018). Depending on the type of stimuli, microglia are capable of acquiring both cytotoxic role and exacerbate brain damage as well as immune regulatory, promoting injury resolution and tissue regeneration (Colonna & Butovsky, 2017; Li & Barres, 2018; Wolf et al., 2017). Microglia quickly reacts to the cues they receive from the neurodegenerative milieu, importantly from neurons and astrocytes. Activated microglia can trigger further astrocytic response and neuroinflammatory processes to contain pathology and promote healing via tissue recovery and regeneration. Therefore, impairment of normal microglial functions or prolonged activation also affects normal brain functioning.

Also astrocytes can directly modulate neuronal function and visibly affect animal behavior (Barnett et al., 2023; Nagai et al., 2021; Purushotham & Buskila, 2023). Astrocytes are mostly regarded as mediators of molecular homeostasis. They provide neuron metabolic, ionic and trophic support, modulate synaptic function, and maintain the blood – brain barrier (BBB) (Fisher & Liddelow, 2024; Tremblay et al., 2019). Similar as microglia, but to a lesser extent and different specification, they are also effectors and propagators of immune signaling with their own repertoire of receptors and signaling molecules, enabling them to detect and respond to inflammatory stimuli (Fisher & Liddelow, 2024). Astrocytes become activated to the spectrum of different phenotypes, both protective and cytotoxic (Escartin et al., 2021). They can activate microglia and peripheral immune cells and recruit them to the site of injury by releasing ATP. Astrocytes attenuate microglial inflammatory responses through the release of GABA and can trigger antioxidant response in microglia. Both glial cell types cooperate to eliminate apoptotic cells and co-ordinately regulate neuronal synaptic pruning (Fisher & Liddelow, 2024; Garland et al., 2022). This bidirectional interaction is a key factor for neuron functioning. Any deviations can induce pathogenesis.

Here, we focused on neurodegenerative diseases using animal model of early Parkinson’s disease by selectively damaging a portion of dopaminergic neurons using 6-OHDA and observing processes involved in response to this damage, particularly driven by astrocytes and microglia. Selective lesion of nigrostriatal system up to the certain threshold causes no or only temporal movement disorder. Reversal of motor dysfunction is known as spontaneous functional compensation, similar as at the early stages of PD. In the previous studies (K. Kuter et al., 2018, 2019) death of astrocytes induced by chronic infusion of fluorocitrate (FC) into the substantia nigra (SN) caused massive microglia activation and stressed neurons. As a result, spontaneous compensation of motor deficits caused by partial dopaminergic neuron loss was blocked. This indicated that astrocytic support was essential for the functional compensation. Nevertheless, concomitant astrocyte loss and microglia activation did not allow to distinguish whether functional disability was due to astrocyte loss or rather strong microglia activation. **In order to separate effect of loss of astrocyte function from the dysfunction caused by activated microglia we used minocycline to inhibit microglia response. We checked how microglia activation inhibition affected neuron and astrocyte cell death caused by selective toxins and locomotor function and potential for spontaneous functional compensation of deficits.**

Antibiotic minocycline is a tetracycline analogue and a potent, though non-specific inhibitor of microglial activation. It suppresses secretion of pro-inflammatory mediators, nitric oxide and chemotaxis of neutrophils, inhibits apoptotic death through caspase-dependent and -independent pathways (Awogbindin et al., 2020). It was also shown that minocycline acted protective to dopaminergic neurons in various animal models of PD (Cronin & Grealy, 2017; Du et al., 2001; Sun et al., 2019; Wang et al., 2020; Wu et al., 2002). But there were also unequivocal (Quintero et al., 2006) and negative reports showing that minocycline was not neuroprotective (Sriram et al., 2006) or even enhanced neurotoxicity (Diguet et al., 2004; Yang et al., 2003). Although, minocycline is listed among the putative neuroprotective agents by the Committee to Identify Neuroprotective Agents in Parkinson’s (CINAPS) in a 12-month clinical trial in early Parkinson’s disease patients (NET-PD FS-1) (Ravina et al., 2006) and its 6 months follow-up NET-PD FS-2 study (Kieburtz K et al., 2008) it did not show any beneficial effect. **Therefore, establishing whether inhibition of microglia activation by minocycline is responsible for neuroprotective and functionally relevant behavioral effect is essential for providing more effective therapies against PD.**

We show in this study that minocycline treatment by decreasing strong microglia activation induced by stress signals from FC-treated astrocytes probably protected portion of astrocytes in the SN. On the other hand, minocycline negatively affected astrocytes in 6-OHDA lesion group where microglia activation was mediated by progressive neuron loss. Therefore, we conclude, that phenotype of microglia activation is different when induced by astrocyte death vs by neuron death. Thus, effects of its inhibition can be either protective or detrimental, respectively. Especially that minocycline treatment induced non-dopaminergic neurodegeneration in SN and ventral tegmental area (VTA). Despite it minocycline improved compensatory potential after combined FC and 6-OHDA treatment. It suggests better functional coupling of remaining astrocytes and neurons in the SN when less microglia was excessively activated. Importantly, those experiments provide evidence for **astrocytes to be a possible targets for non-neuronal treatment of PD-relevant motor behavior dysfunctions.**

## 2. Materials and methods

### 2.1. Animals and drugs

Three months old male Wistar HAN rats (Charles Rivers, Germany) were kept under 12 hour dark/light cycle (light from 06:00 to 18:00), with free access to food and water. The experiments were carried out in compliance with the Animal Experiments Bill of January 21, 2005; (published in Journal of Laws no. 33/2005 item 289, Poland), and according to the EU Directive 2010/63/EU for animal experiments; as well as National Institutes of Health guide for the care and use of laboratory animals (NIH Publications No. 8023, revised 1978). They also received approval from the Local Ethics Committee (approval no: 221/2018; 221A/2018, 306/2018; 33/2020; 242/2020, 259/2022). All efforts were made to minimize the number of animals and their suffering.

Minocycline hydrochloride (Biorbyt LLC. #orb341890, dissolved in 4 ml sterile water with ca. 0,5% 5M NaOH) was administered in a dose 25 mg/kg ip, twice a day for a total of 9 days, with surgery on day 2, behavioral testing on days 5 and 8 (3 and 6 days after surgery, respectively) and sacrifice on day 9.

### 2.3. Brain surgery

Stereotaxic brain operations were performed according to Kuter *et al*. (2016), under ketamine and xylasine anesthesia (65 - 50 mg/kg and 10 - 3 mg/kg i*m*, Biowet, Puławy, Poland). Desipramine (30 mg/kg *ip*, Sigma-Aldrich, Germany) was administered 30 min before lesioning to protect the noradrenergic terminals. To induce degeneration of dopaminergic neurons the animals were stereotaxically, bilaterally injected with 6-OHDA HBr (3 μg base/3 μl per side), dissolved in 0.2% ascorbic acid (both from Sigma-Aldrich, Germany) into the passing fibers of the medial forebrain bundle (MFB), at coordinates: AP: 1.4 mm, L: ± 1.6 mm, V: 8.7 mm from bregma, according to Paxinos and Watson’s atlas (2007). Control - sham operated rats received solvent in the same way. The injection cannula was left in place for 2 min for full absorption of the solution. Additionally, in the same animals, stainless steel cannulas were bilaterally, permanently implanted in the SN *pars compacta* (SNc) (coordinates: AP: 4.9 mm, L: ± 1.8 mm, V: 8.3 mm from bregma, according to Paxinos and Watson’s atlas (2007)) and connected by a catheter to osmotic minipumps (1007D, ALZET, Charles-Rivers, Germany), implanted under skin on the neck, that administered fluorocitrate (FC, 2 nmol/day, Sigma-Aldrich, Germany) for 7 days, at a continuous rate 0.5 μl / hour, to induce astrocyte dysfunction. Respective control animals had cannulas implanted with sealed catheters. FC was prepared according to (Paulsen et al., 1987). The rats received 1 ml of 20% glucose solution (Pol-Aura, Poland) and meloxicam (1 mg/kg, sc, Bioveta, Czechia) on the day of operation and 24 hr afterwards. Body weight of animals was monitored during the whole experiment.

The animal groups were as followed: SSS – sham / sham / solvent; SFS - sham / FC / solvent; LSS – 6-OHDA lesion / sham / solvent; LFS – 6-OHDA lesion / FC / solvent; SSM – sham / sham / minocycline; SFM - sham / FC / minocycline; LSM – 6-OHDA lesion / sham / minocycline; LFM – 6-OHDA lesion / FC / minocycline. Number of animals per group was 5 - 9. Each animal is represented on the graph as a single dot.

### 2.3. Behavioral analysis of locomotor activity

Rat locomotor activity (path length, locomotion and resting times) and rearings (total, free and supported, number, duration) were measured at two time-points after operation (3^rd^ and 6^th^ days) using computerized actimeters (ACTIFRAME-SYSTEM, GERB Elektronik GmbH, Berlin; Germany, with ARNO software) as described before in detail (Kolasiewicz et al., 2012; K. Kuter et al., 2016). Animals were placed in the cages individually with free access to food and water and their behavior was analyzed for 60 minutes.

### 2.4. HPLC analysis of dopamine and its metabolites

Rats were decapitated on the 7^th^ day after operation. Single STR and SN were immediately dissected and frozen on dry ice. Tissue was kept at –80°C until further analysis. The levels of DA and its metabolites: 3,4-dihydroxyphenylacetic acid (DOPAC), 3-methoxytyramine (3-MT), homovanillic acid (HVA) were assessed using HPLC method with electrochemical detection as described previously (Berghauzen-Maciejewska et al., 2014). Briefly, tissue samples were homogenized in 0.1 M perchloric acid containing 0.05 mM ascorbic acid and injected into the HPLC system (column Hypersil Gold C18, 100 x 3.0 mm, 3 μm, Thermo Scientific, UK) equipped with electrochemical detector analytic cell 5010 Coulochem III (ESA, Inc. USA). The mobile phase was composed of 50 mM NaH_2_PO_4_ x 2H_2_O; 40 mM citric acid; 0.25 mM 1-octanesulfonic acid sodium salt; 0.25 mM EDTA; 1.3% acetonitrile; 2.4% methanol. The applied potential was E1 = –175 mV and E2 = +350 mV. The data were quantified using the area under peaks and external standards with Chromeleon software (Dionex, Germany). The turnover rates were calculated as metabolites to neurotransmitter ratios.

### 2.5. Immunohistochemistry

After decapitation the brain hemisphere was rapidly removed, postfixed in the cold 4% paraformaldehyde and cryoprotected in 20% sucrose solution. The brains were cut on a freezing microtome into 25 μm frontal sections (AP –4.4 to 6.6 mm from bregma according to Paxinos and Watson 2007 for SNc - VTA) and stained as described before (Kuter et al., 2011).

For tyrosine hydroxylase (TH) and Nissl, S100 and Iba-1 staining sections were incubated in primary antibodies (anti–TH; AB_2201526 from Chemicon Int., USA; anti-Iba-1 AB_839504; WAKO, Japan). For anti-S100 (AB_306716; Abcam, UK) staining, heat induced antigen retrieval in 10mM citrate buffer, pH 6.0 was performed. After incubation with secondary antibodies (anti-mouse (AB_2313571) or anti-rabbit (AB_2313606) biotinylated, Vector Laboratories, UK) sections were processed using an ABC-peroxidase kit (Vector Laboratories, UK) and 3,3’-diaminobenzidine as a chromogen. Subsequently, sections containing SNc – VTA stained for TH were counterstained with 1% cresyl violet (CV) with Nissl method. All sections were cover-slipped in a Permount medium (Fisher Scientific, USA).

### 2.6. Morphological analysis of microglia – Sholl analysis

Iba-1^+^ cells were photographed under 100x magnification at the systematic, random positions chosen by a newCAST (Visiopharm, Denmark) software with a light microscope (Leica, Denmark). Per each rat approx. 22 cells were counted on two tissue sections close to the injection cannula site in the SNc. Analysis was performed using Fiji ImageJ software. The scale was converted from pixels to micrometers. Each image was split into single channels, turned into grayscale, turned into binary image using standard thresholding. Selected cells with all branches visible in the image frame were separated, followed by manual selection of the cells’ centre of mass. Single cell analysis was carried out using the Sholl analysis built into Fiji, which was performed with parameters: starting radius:10 μm; ending radius: 50 μm; radius step size: 10 μm. Number of cell branches crossing each radius were counted. Ramification index (Schoenen’s factor) - ratio between the maximum number of intersections with a circle and the number of primary protrusions was analysed for each cell.

### 2.7. Cell counting in the SNc

TH^+^ and/or CV^+^ neurons, S100^+^ astrocytes and Iba-1^+^ microglia cells were counted in the SNc and VTA as described previously (K. Kuter et al., 2007). Stereological counting was performed using a light microscope (Leica, Denmark) controlled by a newCAST (Visiopharm, Denmark) software. The analyzed regions were outlined under lower magnification (5x) and their areas were estimated. The number of stained cells was calculated under 100x magnification using a randomized meander sampling and the optical dissector methods.

### 2.8. Statistics

Results are presented as the mean ± standard error of mean (SEM). The statistical analysis of results was performed using STATISTICA 10.0 software (StatSoft Inc., USA) or GraphPad Prism 10 (GraphPad Software, LLC, Boston, MA, USA). P ≤ 0.05 was considered as statistically significant and 0.1

≥ p > 0.05 were considered as trends. Analyses were done by two way ANOVA with the Fisher Least Significant Difference (LSD) *post hoc* test and t test for comparison of groups in time. A repeated measures ANOVA test was used for motor behavior analysis in different time-points.

## 3. Results

### 3.1. Verification of the astrocyte death by FC

Prolonged FC infusion caused death of S100b^+^ astrocytes in the SNc (SSS = 8048, SFS = 4802 cells/mm^3^, t=4,474, df=10, p=0,0012, fig. 3). The size of this effect was the same in both groups FC alone (SFS) and in FC accompanied by 6-OHDA injection (LFS = 4764 cells/mm^3^, t=3,912, df=13, p=0,0018). 6-OHDA neuronal lesion of SNc did not affect astrocyte survival. Minocycline treatment decreasing microglia activation partially prevented astrocytes from death caused by FC, as shown by lack of significant differences between control SSM vs SFM and LFM groups. This result needs to be further verified because there was no significant difference between solvent-treated vs minocycline-treated FC groups (SFS vs SFM). Minocycline lowered density of astrocytes in 6-OHDA group, as compared between solvent and minocycline treatment (LSS vs LSM, t=2,933, df=13, p=0,0117). Minocycline had no significant influence on combined FC and 6-OHDA treatment.

**Fig. 1.**
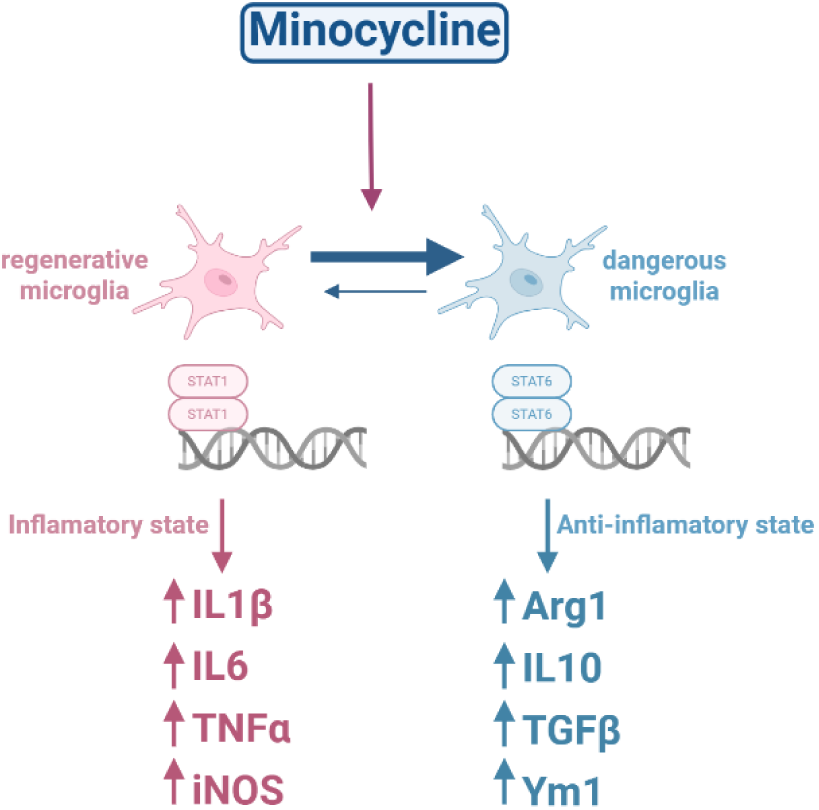
Schematic summary of minocycline mechanism of anti-inflammatory action, inhibiting activation of microglial state and shift in production of cytokines and inflammation markers.

**Fig. 2.**
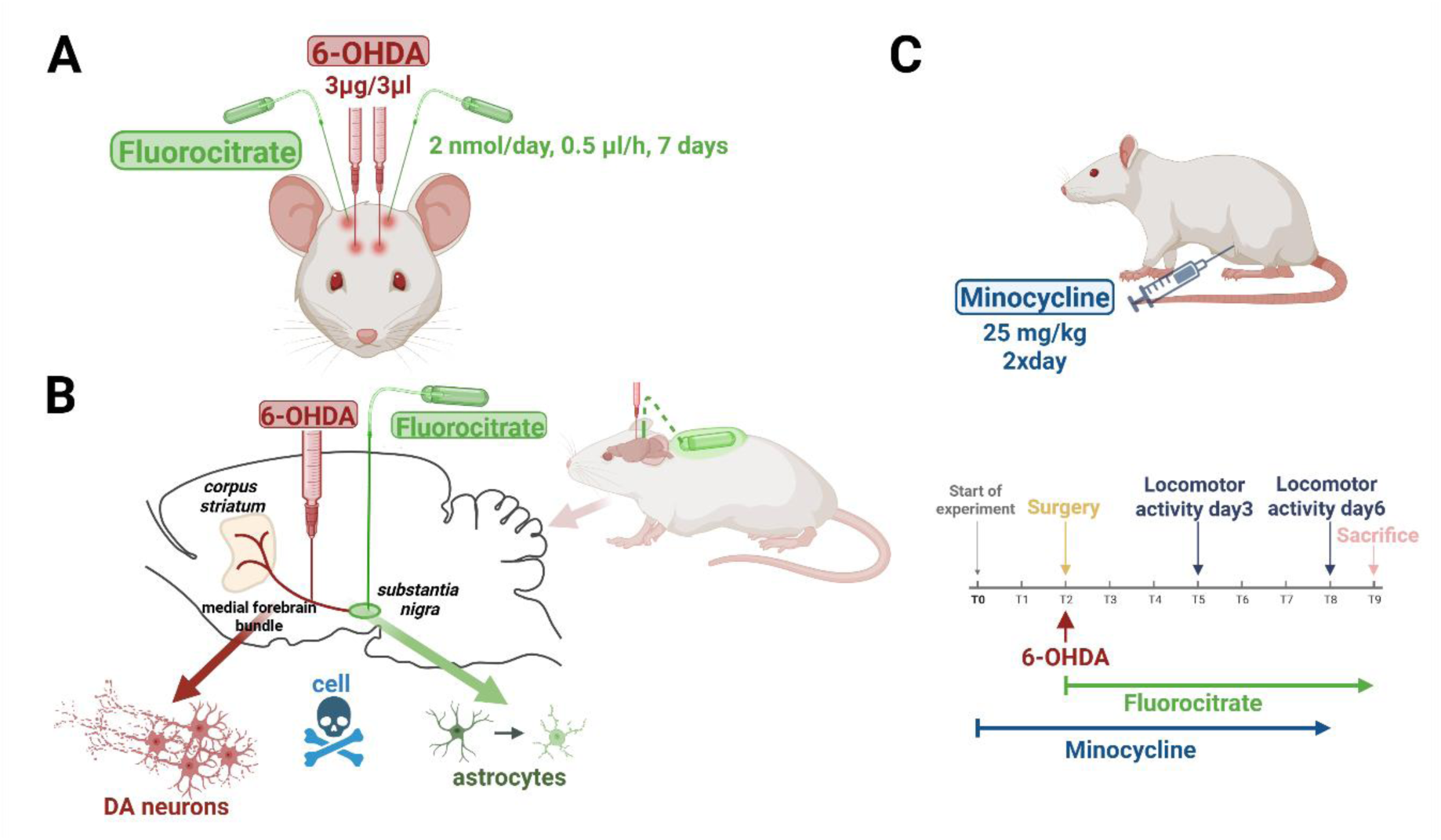
Schematic representation of experimental plan and timeline of central toxin (fluorocitrate, FC and 6-OHDA) injection and peripheral minocycline/solvent administration.

**Fig. 3.**
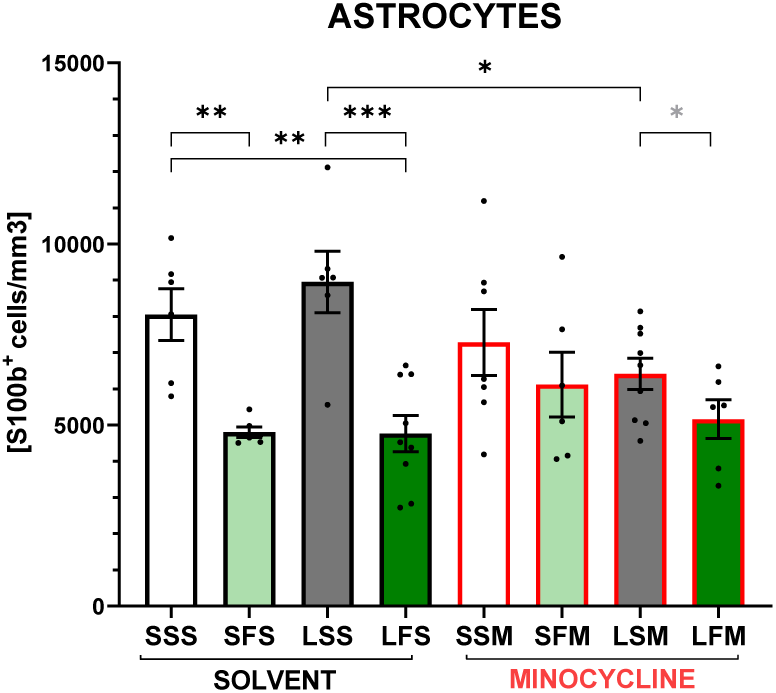
The stereological counting of astrocytes (S100b^+^) in substantia nigra pars compacta (SNc). Results are shown as mean ± SEM cell number per mm^3^. Each dot represents a value from a single animal. In the group names S stands for sham or solvent, F – fluorocitrate, L – 6-OHDA lesion, M – minocycline. Two way ANOVA with Fisher Least Significant Difference post hoc test was used with p ≤ 0.05 marked as significant (*). 0.1≥p ≥0.05 were considered as trends and marked in grey colour.

### 3.2. Morphological verification of the microglia activation inhibition by minocycline

Chronic FC infusion caused severe microglia activation as a reaction to astrocyte dysfunction and death in the SNc. This was documented here by morphological analysis of Iba-1^+^ cells. Both ramification index and mean number of ramifications showed significantly reduced arborisation of microglial cells shape due to FC (fig.4). Minocycline diminished this morphological change in both FC groups. 6-OHDA did not change ramification index but decreased mean number of intersections (LSS vs SSS, p= 0,0085) showing only subtle influence of neuron death on microglia activation state. Interestingly, microglial cells in FC combined with 6-OHDA-treated group were less affected morphologically than FC alone group and minocycline was not effective there.

**Fig. 4.**
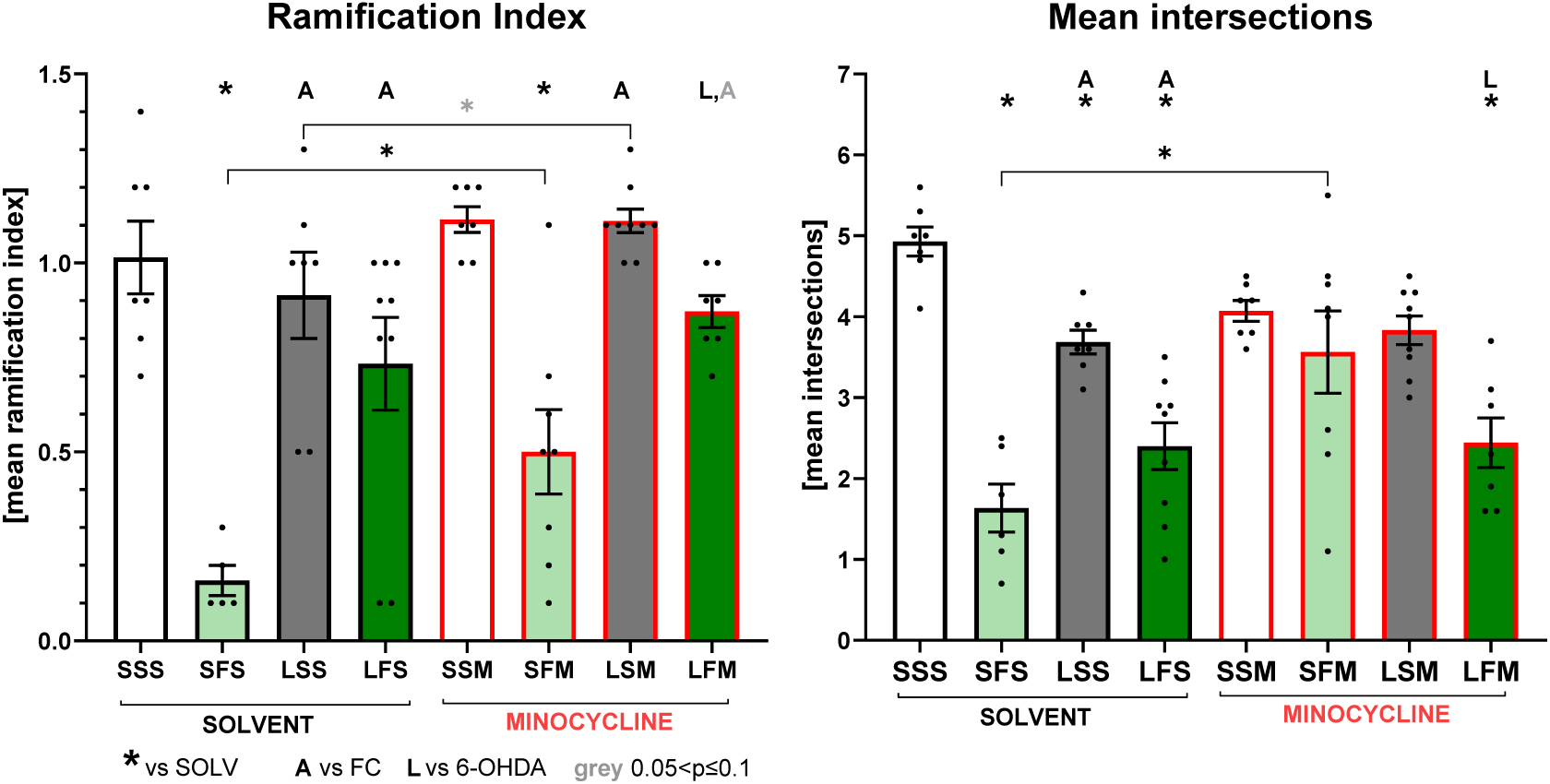
Ramification index and mean number of ramifications of microglia in substantia nigra pars compacta (SNc). Results are shown as mean ± SEM. Each dot represents a mean value from multiple microphotographed cells from a single animal. In the group names S stands for sham or solvent, F – fluorocitrate, L – 6-OHDA lesion, M – minocycline. Two way ANOVA with Fisher Least Significant Difference post hoc test was used with p ≤ 0.05 marked as significant vs solvent (*), vs minocycline (M), vs FC (A), vs 6-OHDA (L). 0.1≥p ≥0.05 were considered as trends and marked in grey colour.

### 3.3. Verification of the neuronal lesion size caused by 6-OHDA

6-OHDA lesion caused selective degeneration of dopaminergic neurons in SNc (SSS = 10237, LSS = 7479, p = 0,0209) (fig. 5), non-dopaminergic neurons were not affected significantly. VTA dopaminergic neurons were also not affected. FC decreased neuron density in SNc (SFS = 8453, p = 0,0835 for dopaminergic neurons and SSS = 6650, SFS = 5178, p=0,0334 for non-dopaminergic neurons), but not in VTA. Combined treatment with both toxins did not aggravate significantly neurodegeneration.

**Fig. 5.**
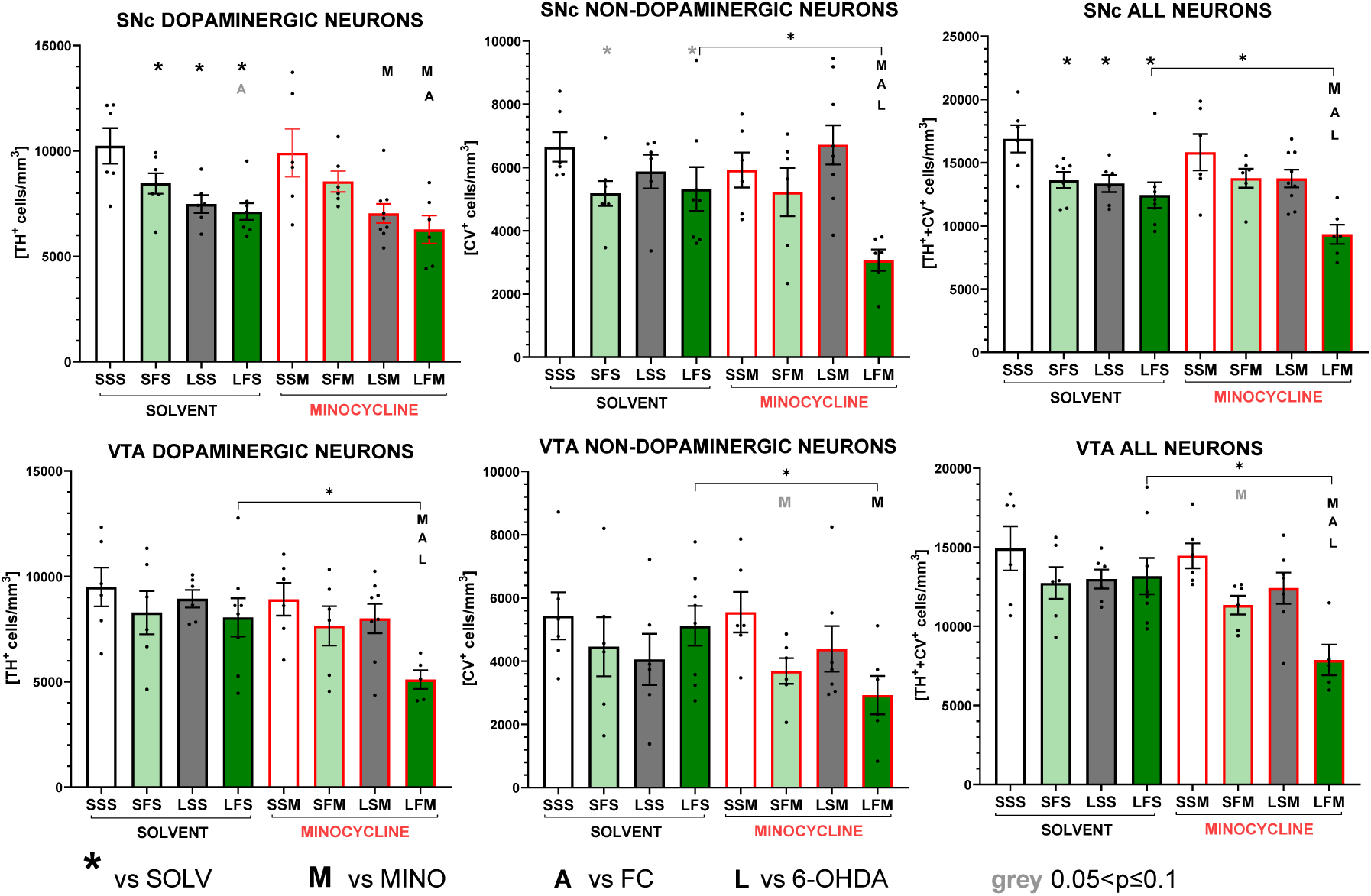
The stereological counting of dopaminergic (TH^+^, A, D) and non-dopaminergic (cresyl violet only stained, CV^+^, B, E) neurons in substantia nigra pars compacta (SNc, A – C) and ventral tegmental area (VTA, D – F). C and F show values both neuron types. Results are shown as mean ± SEM cell number per mm^3^. Each dot represents a value from a single animal. In the group names S stands for sham or solvent, F – fluorocitrate, L – 6-OHDA lesion, M – minocycline. Two way ANOVA with Fisher Least Significant Difference post hoc test was used with p ≤ 0.05 marked as significant vs solvent (*), vs minocycline (M), vs FC (A), vs 6-OHDA (L). 0.1≥p ≥0.05 were considered as trends and marked in grey colour.

Chronic administration of minocycline itself or combined with FC did not affect neuron density in midbrain. Non-significant shift from neurons with dopaminergic markers (TH^+^) to non-dopaminergic (CV^+^) was visible in LSM group indicating loss of dopaminergic phenotype but not cell death, as discussed previously (K. Kuter et al., 2016). Overall neuronal density was not changed in this group. Minocycline induced neurodegeneration after combined 6-OHDA and FC treatment in the SNc affecting mostly non-dopaminergic neurons (LFS = 5321, LFM = 3069, p = 0,0154 for non-dopaminergic neurons) and both types of neurons in the VTA (LFS = 8056, LFM = 5109, p = 0,0159 for dopaminergic neurons and LFS = 5118, LFM = 2927, p = 0,0279 for non-dopaminergic neurons).

Dopamine (and its metabolites DOPAC, HVA, 3-MT – not shown) level in the STR tissue homogenates was strongly decreased after dopaminergic neuron death in the SNc (fig. 6). DA turnover rate was significantly elevated. Neither FC treatment, astrocyte death and microglia activation, nor minocycline anti-inflammatory treatment had significant influence on both parameters.

**Fig. 6.**
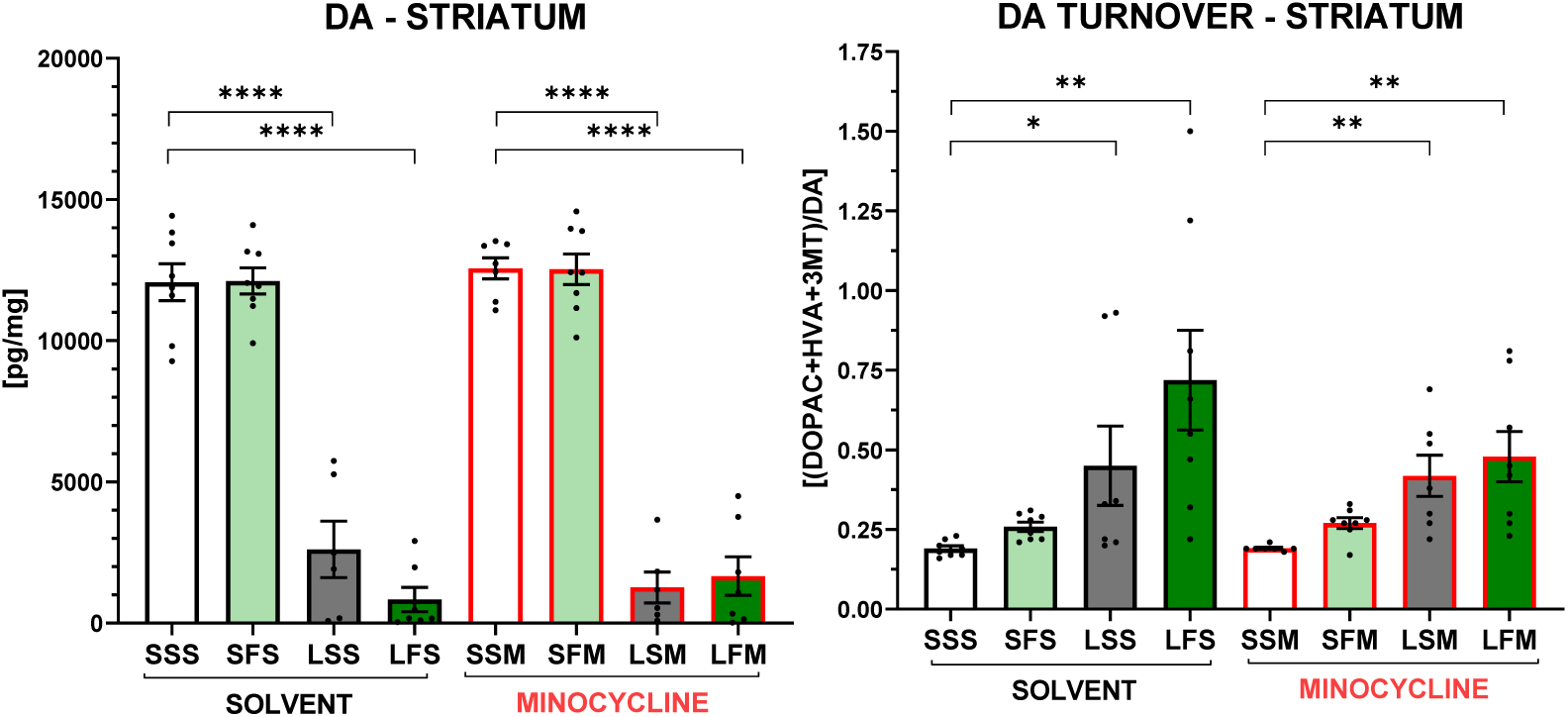
HPLC analysis in striatum of dopamine (DA), and turnover rate calculated as sum of its metabolites (3,4-dihydroxyphenylacetic acid (DOPAC), 3-methoxytyramine (3-MT), homovanillic acid (HVA)) divided by DA level. Results are shown as mean ± SEM. Each dot represents a value from a single animal. In the group names S stands for sham or solvent, F – fluorocitrate, L – 6-OHDA lesion, M – minocycline. Two way ANOVA with Fisher Least Significant Difference post hoc test was used with p ≤ 0.05 marked as significant vs solvent (*).

### 3.4. Behavioral markers of locomotor deficit functional compensation

Locomotor activity was measured to validate selective dopaminergic neurodegeneration and induction of motor deficits, similar as in Parkinson’s disease. In early PD rat model of medium size dopaminergic neuron lesion locomotor disability spontaneously recovers within 1 - 2 weeks as shown previously (K. Kuter et al., 2016, 2018). At the first week hyperactivation is usually observed. Comparison of path length data between the 3^rd^ and 6^th^ day after operation was analysed as a measure of spontaneous functional motor compensation in each rat. 6-OHDA injection into MFB caused visible decrease in rat locomotion 3 days after injection and its recovery after 6 days showing a significant potential to functionally compensate motor deficits (fig. 7). FC treatment ablated such recovery but minocycline treatment recovered it. Minocycline improved compensatory potential in sham operated animals (SSM vs SSS p=<0,0001) and in combined toxin group (LFM vs LFS p=0,0182) but not after FC alone. Visible minocycline-induced decrease in compensatory potential after 6-OHDA alone did not reach statistical significance (p=0,106).

**Fig. 7.**
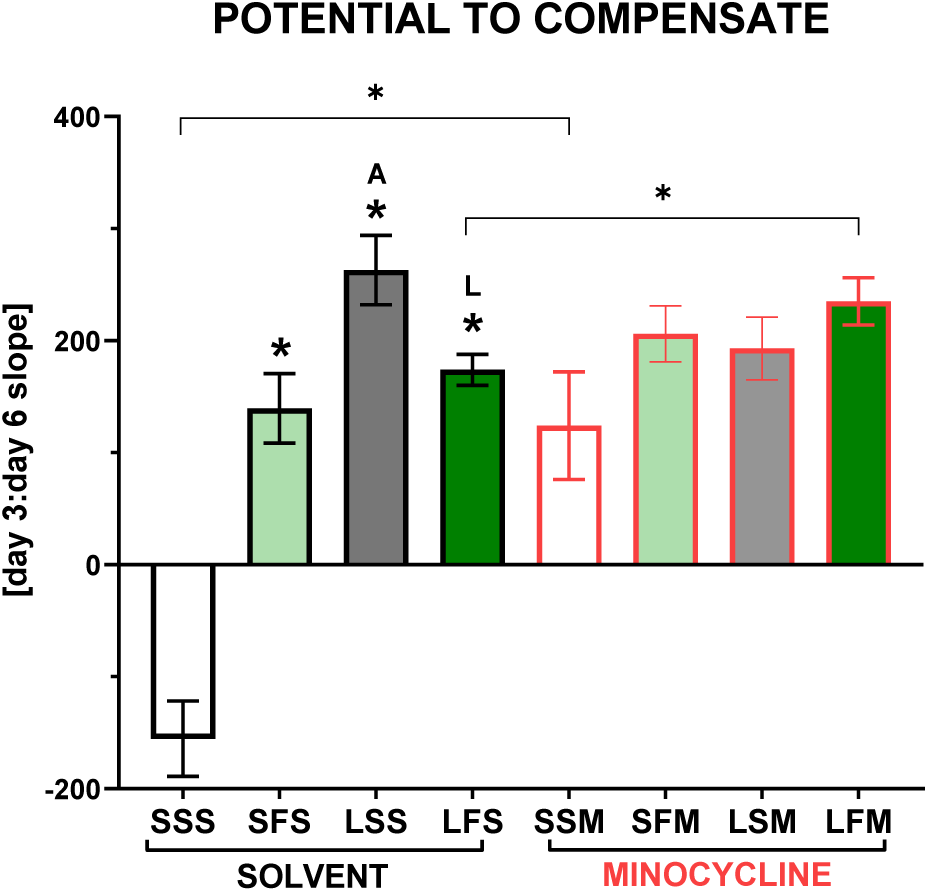
Functional potential to compensate against locomotor dysfunction was measured as a slope of a line drawn between points representing value of path length walked by animal between third and sixth day of behavioral testing after operation. Results are shown as mean ± SEM. In the group names S stands for sham or solvent, F – fluorocitrate, L – 6-OHDA lesion, M – minocycline. Two way ANOVA with Fisher Least Significant Difference post hoc test was used with p ≤ 0.05 marked as significant vs solvent (*), vs FC (A), vs 6-OHDA (L).

## 4. Discussion

Early research on inflammation in PD indicated that inhibition of microglia activation was neuroprotective. This was true mostly when strong inflammation was induced as for example hemorrhagic or ischemic stroke, acute injury or with LPS (Elewa et al., 2006; Tomás-Camardiel et al., 2004). But with further development on physiological role of active microglia in regeneration and neuronal functioning adaptations brought understanding that active microglia has also its important role and more beneficial should rather be shifting microglia activation towards more regenerative state than freezing activation at all.

### Differential microglia activation due to astrocyte vs neuronal death

In this study, death of approx. 40% astrocytes induced by chronic infusion of FC into SNc strongly activated microglia, observed as reduced arborisation of microglial cells (decreased both ramification index and mean number of ramifications). This is in line with our previous studies (K. Kuter et al., 2018) in this model showing reduced density of S100^+^ astrocyte cell bodies in the SN and in consequence massive activation of microglial cells visualized with Iba-1 in tissue sections and in Western blot. On the other hand, we found here that loss of 27% dopaminergic neurons caused only minor activation of microglia visible as decreased mean number of ramifications but not changed ramification index (Schoenen’s factor). This further confirms previous observation of immunohistochemical staining with Iba-1 in the SNc where microglial cell activation due to 33 % loss of dopaminergic neurons was manifested by a stronger microglia staining but with terminals still long and branched, only slightly thicker and with varicosities. In contrast to FC effect, Iba-1 protein amount was not changed (K. Kuter et al., 2018). Observed difference in Schoenen’s factor between FC-activated vs 6-OHDA-activated microglia suggests that **dying astrocytes induce different phenotype of microglia activation than degenerating neurons**. Of course, it needs to be taken into account that FC was infused for 7 days directly into analyzed SNc, while 6-OHDA was acutely injected into MFB and neurons degenerated slowly by dying-back process starting with their terminals and progressing towards cell bodies. Either way both cell type deaths were progressive.

### Astrocyte loss affected neurons but neuron loss did not affect astrocytes

In the previous studies (K. Kuter et al., 2018) decreased density of astrocytes after FC treatment reached 26%. The conclusion from previous studies was that FC stressed but did not kill neurons itself. But when combined with 6-OHDA accelerated neuronal cell death. Similar results were obtained using L-Aminoadipic Acid (L-AA) to kill astrocytes in the striatum without affecting nerve terminals (Voronkov et al., 2021). Here, astrocyte loss was larger, up to 40% in the SNc. This could be the reason that neuronal death was documented in consequence (both dopaminergic and non-dopaminergic), and no enhanced effect was observed 7 days after operation, after combined FC and 6-OHDA treatment. It means that **there is a threshold between 26 and 40% of astrocyte loss that after crossing can be deleterious for neurons**.

When looking at the opposite direction of interaction, namely how neuronal cell death affects astrocytes we did not observe any changes in the SN due to 6-OHDA injection, similar as in the previous study analyzing GFAP, S100 and ALDH1L1 markers (Kuter et al., 2018).

### Microglia activation profile is opposite when induced by astrocyte vs neuronal death

Microglial cells in FC combined with 6-OHDA-treated group were less affected morphologically than FC alone group. It was visible especially well in Schoenen’s factor where ramification index was significantly higher than in FC alone group. This suggests that **astrocyte death activates microglia by not only different but probably opposite mechanism than neuronal death**.

### Minocycline treatment decreased morphological signs of microglia activation and protected some astrocytes

In order to distinguish the effect of astrocyte death form microglia activation minocycline treatment was used. It diminished morphological signs of microglia activation visible as reduced arborisation of microglial cells due to FC. Our data indicate that FC-induced astrocyte death is mostly caused directly by FC inhibition of aconitase, but has also a minor part that is driven by strongly activated microglia which exacerbates astrocyte death. Decreased activation of microglia by minocycline prevented cell death of a portion of astrocytes. This partial inhibition of strong microglia activation by minocycline could be the reason for smaller size of astrocyte death in the SN as compared to FC alone. The conclusion of astrocyte protection by minocycline needs to be further verified. We based it on the lack of significant difference between sham operated and FC-treated animals, both receiving minocycline, while solvent treated rats showed significant loss of astrocytes (see fig. 3). It is known that strong activation of microglia can be toxic to bystander cells (Butler et al., 2021; Pampuscenko et al., 2025). The astrocyte death could be driven not only by chronic and selective aconitase inhibition by FC but also by microglia toxicity. Inhibition of part of microglia could have **protected portion of astrocytes from inflammation-driven degeneration**.

### Could minocycline treatment be toxic towards astrocytes interacting with degenerating neurons?

Chronic minocycline treatment significantly lowered density of S100^+^ astrocytes in the SNc of 6-OHDA treated group, when compared data between solvent and minocycline treatment. But when to compare minocycline treatment alone vs with 6-OHDA (SSM vs LSM) there was no significant decrease. This observation was mostly due to the dispersion of data in the SSM control group. No influence of microglia inhibition on astrocytes after combined FC and 6-OHDA treatment stands against such astro-toxic mechanism. In the previous study by Stefanova et al. 4 weeks of minocycline treatment inhibited both microglia and astrocytes (Stefanova et al., 2004). They showed decreased GFAP density in the striatum which could suggest also loss of astrocytes. Here minocycline was administered for much shorter time, twice a day for 9 days. It needs to be further verified whether prolonged minocycline treatment could be potentially dangerous for astrocytes. Further analysis of astrocyte activation markers and morphology in response to minocycline could be helpful in this manner. As discussed above, using several protein markers we did not identify any changes in astrocytes in response to neurodegeneration (K. Kuter et al., 2018). But there could be other markers, more specific for subtle phenotype changes that would show what happens to astrocytes due to neuron death and how minocycline potentially affects this relation.

### Was minocycline treatment neuroprotective?

Previous data has shown differential effects of minocycline towards neurons – as neuroprotective (mutant alpha-synuclein overexpression) (Wang et al., 2020),6-OHDA Wistar rat striatum (Landim Falcão Tavares Ferreira & De Barros Viana, 2015), 6-OHDA mouse striatum (He et al., 2001), Zebrafish 6-OHDA (Cronin & Grealy, 2017), rotenone (Sun et al., 2019), MPTP in mice (Du et al., 2001; Wu et al., 2002), unequivocal (Quintero et al., 2006), not neuroprotective (6-OHDA to MFB and quinolinic acid to striatum (Stefanova et al., 2004), MPTP models (Sriram et al., 2006) or enhanced neurotoxicity (MPTP non-human primates) (Diguet et al., 2004; Yang et al., 2003). In this study we show that minocycline had no influence on dopaminergic neuron death induced by 6-OHDA but negatively affected non-dopaminergic neurons after combined 6-OHDA and FC treatment in the SNc and VTA. 6-OHDA injected in 3 µg dose to MFB under desipramine blockade of noradrenergic transporter was proven to be selective towards dopaminergic neurons (K. Kuter et al., 2018). Our standard procedure always quantifies both dopaminergic and non-dopaminergic neurons at the same tissue sections to verify actual neuronal degeneration rather than loss of TH and dopaminergic phenotype as we described before (K. Kuter et al., 2016). In a similar paradigm study Du et al. also quantified TH^+^ SNc neurons stereologically and estimated dopamine and DOPAC levels in the striatum after 7 day minocycline treatment. Instead of rat 6-OHDA they used male C57BL6 mice MPTP model and have proven neuroprotection towards dopaminergic cells (Du et al., 2001). But they did not quantify non-dopaminergic neurons, therefore potential minocycline toxicity towards non-dopaminergic neurons could have been missed. Also, MPTP has more pronounced pro-inflammatory effect, as glial cells are essential for MPTP metabolism to toxic MPP+ and microgliosis is stronger than in 6-OHDA models, especially injected to MFB, where microglia activation is driven purely by neuron death. Other studies concluded neuroprotection indirectly, based on behavioral output or dopamine levels (Landim Falcão Tavares Ferreira & De Barros Viana, 2015; Stefanova et al., 2004). Other studies failed to confirm neuroprotection or even reported deleterious effects of minocycline on neuronal survival in PD mouse models (Yang et al., 2003). Why non-dopaminergic neurons are more vulnerable in SNc than dopaminergic as we showed here needs to be further elucidated. Another issue worth taking under consideration but not studied here is the potentially negative effect of prolonged antibiotic administration on gut microbiota (Markulin et al., 2022).

### Minocycline improved behavioral outcome despite lack of neuroprotection

Similar as patients at the early stages of PD, who do not show motor dysfunction despite progressive neurodegeneration, medium size lesion of the rat nigrostriatal system is easily compensated functionally and motor dysfunction is spontaneously reversed with time (Bezard et al., 2003; Bezard & Gross, 1998; Blesa et al., 2017; K. Kuter et al., 2016, 2018, 2019; K. Z. Kuter et al., 2021; Ossowska et al., 2005; T. Robinson et al., 1994; T. E. Robinson & Whishaw, 1988)). This and previous studies have shown that FC-driven loss of astrocytes and microglia activation blocked spontaneous compensation of locomotor deficits (K. Kuter et al., 2018). Interestingly, minocycline treatment did not rescue dopaminergic neurons in SNc, nor dopamine levels and turnover in the striatum, even caused non-dopaminergic neuron loss, but still enhanced compensatory potential to functionally improve walking after combined 6-OHDA lesion and FC-induced astrocyte death. Taking into account that the changes were observed within the first week post lesion it suggests that at this stage of medium size nigrostriatal system lesion these are still rather dopaminergic than other types of neurons in SN that are responsible for compensation. Similarly in the work of Voronkov et al. (Voronkov et al., 2021) local administration of gliotoxin L-AA in the striatum of rats caused astrocytic degeneration without affecting the neurons and nigrostriatal fibers but the failure of astrocyte-neuron coupling in the striatum lead to motor dysfunction such as gait disturbances and bradykinesia. Therefore it is possible that minocycline by decreasing microglia activation state helped to at least partially maintain such astrocyte-neuron coupling. At this stage it is not possible to clearly interpret those data and the answer might lay in other parameters not studied here. Recent study by Evans et al. 2025 established an important, novel functional role of striatal astrocyte signalling in modulating motor function in PD. They have proven that motor impairment in dopamine-depleted animals can be improved by manipulation with astrocytes (Evans et al., 2024). Microglia function is essential in neuron adaptation by rebuilding synapses together with astrocytes. This could also be the case in FC and 6-OHDA treated animals that stronger motor compensation due to minocycline was rather an outcome of better functional coupling of remaining astrocytes and neurons in the SN when less microglia was overly activated. Importantly, those experiments together provide evidence for **astrocytes as an important contributors to treatment of PD-relevant motor behavior dysfunctions. The main advantage of non-neuronal target for potential PD therapeutics is that astrocytes can repopulate after partial degeneration**.

### Conclusions

We provide a note of caution regarding the success of microglial-targeted PD strategies, using minocycline as an example. Pure inhibition of inflammation and glial activation is not the best neuroprotective approach in slowly progressing neurodegenerative diseases as activated microglia is necessary for tissue regeneration. Recent approach to reprograming activated microglia towards more protective phenotype may offer a new strategy to alter the course or mitigate the progression of DA neurodegeneration. But understanding of broader context of microglia – astrocyte – neuron interaction is necessary to obtain effective therapeutic outcome.

## Abbreviations

6-OHDA: 6-hydroxydopamine
FC: fluorocitrate
GFAP: glial fibrillary acidic protein
Iba-1: ionized calcium-binding adaptor molecule 1
MFB: medial forebrain bundle
SN: substantia nigra
SNc: substantia nigra pars compacta
STR: striatum
VTA: ventral tegmental area

## Acknowledgements

The study was supported by the: National Science Centre grant OPUS14 2017/27/B/NZ7/00289 and Statutory Funds of the Maj Institute of Pharmacology, Polish Academy of Sciences in Krakow, Poland. We thank Dominika Biała for technical assistance.

## 6. Authors’ contributions

KZK designed study, applied for funding, supervised the project, performed the experiments, analyzed data, wrote and revised the manuscript. AMJ, MP, JK, BK, IL, WM, TL, JK performed the experiments, analyzed data, JK performed the experiments, prepared figures, edited manuscript.

## 7. Funding

The study was supported by the: National Science Centre grant OPUS14 2017/27/B/NZ7/00289 and statutory funds of the Maj Institute of Pharmacology, Polish Academy of Sciences in Krakow, Poland.

## 8. Declaration of Competing Interest

The authors declare that they have no known competing financial interests or personal relationships that could have appeared to influence the work reported in this paper.

## 9. Data availability

Data will be made available on request to corresponding author.

## Notes

### Competing Interest Statement

The authors have declared no competing interest.

## References

Awogbindin, I. O., Ishola, I. O., St-Pierre, M.-K., Carrier, M., Savage, J. C., Di Paolo, T., & Tremblay, M.-È. (2020). Remodeling microglia to a protective phenotype in Parkinson’s disease? Neuroscience Letters, 735, 135164. 10.1016/j.neulet.2020.135164

Ayerra, L., Abellanas, M. A., Basurco, L., Tamayo, I., Conde, E., Tavira, A., Trigo, A., Vidaurre, C., Vilas, A., San Martin-Uriz, P., Luquin, E., Clavero, P., Mengual, E., Hervás-Stubbs, S., & Aymerich, M. S. (2024). Nigrostriatal degeneration determines dynamics of glial inflammatory and phagocytic activity. Journal of Neuroinflammation, 21(1), 92. 10.1186/s12974-024-03091-x

Barnett, D., Bohmbach, K., Grelot, V., Charlet, A., Dallérac, G., Ju, Y. H., Nagai, J., & Orr, A. G. (2023). Astrocytes as Drivers and Disruptors of Behavior: New Advances in Basic Mechanisms and Therapeutic Targeting. The Journal of Neuroscience, 43(45), 7463–7471. 10.1523/JNEUROSCI.1376-23.2023

Berghauzen-Maciejewska, K., Kuter, K., Kolasiewicz, W., Głowacka, U., Dziubina, A., Ossowska, K., & Wardas, J. (2014). Pramipexole but not imipramine or fluoxetine reverses the “depressive-like” behaviour in a rat model of preclinical stages of Parkinson’s disease. Behavioural Brain Research, 271, 343–353. 10.1016/j.bbr.2014.06.029

Bezard, E., & Gross, C. E. (1998). Compensatory mechanisms in experimental and human Parkinsonism: towards a dynamic approach. Progress in Neurobiology, 55(2), 93–116. 10.1016/S0301-0082(98)00006-9

Bezard, E., Gross, C. E., & Brotchie, J. M. (2003). Presymptomatic compensation in Parkinson’s disease is not dopamine-mediated. Trends in Neurosciences, 26(4), 215–221. 10.1016/S0166-2236(03)00038-9

Blesa, J., Trigo-Damas, I., Dileone, M., del Rey, N. L.-G., Hernandez, L. F., & Obeso, J. A. (2017). Compensatory mechanisms in Parkinson’s disease: Circuits adaptations and role in disease modification. Experimental Neurology, 298, 148–161. 10.1016/j.expneurol.2017.10.002

Butler, C. A., Popescu, A. S., Kitchener, E. J. A., Allendorf, D. H., Puigdellívol, M., & Brown, G. C. (2021). Microglial phagocytosis of neurons in neurodegeneration, and its regulation. Journal of Neurochemistry, 158(3), 621–639. 10.1111/jnc.15327

Colonna, M., & Butovsky, O. (2017). Microglia Function in the Central Nervous System During Health and Neurodegeneration. Annual Review of Immunology, 35(1), 441–468. 10.1146/annurev-immunol-051116-052358

Cronin, A., & Grealy, M. (2017). Neuroprotective and Neuro-restorative Effects of Minocycline and Rasagiline in a Zebrafish 6-Hydroxydopamine Model of Parkinson’s Disease. Neuroscience, 367, 34–46. 10.1016/j.neuroscience.2017.10.018

Diguet, E., Fernagut, P., Wei, X., Du, Y., Rouland, R., Gross, C., Bezard, E., & Tison, F. (2004). Deleterious effects of minocycline in animal models of Parkinson’s disease and Huntington’s disease. European Journal of Neuroscience, 19(12), 3266–3276. 10.1111/j.0953-816X.2004.03372.x

Du, Y., Ma, Z., Lin, S., Dodel, R. C., Gao, F., Bales, K. R., Triarhou, L. C., Chernet, E., Perry, K. W., Nelson, D. L. G., Luecke, S., Phebus, L. A., Bymaster, F. P., & Paul, S. M. (2001). Minocycline prevents nigrostriatal dopaminergic neurodegeneration in the MPTP model of Parkinson’s disease. Proceedings of the National Academy of Sciences, 98(25), 14669–14674. 10.1073/pnas.251341998

Elewa, H. F., Hilali, H., Hess, D. C., Machado, L. S., & Fagan, S. C. (2006). Minocycline for Short-Term Neuroprotection. Pharmacotherapy, 26(4), 515–521. 10.1592/phco.26.4.515

Escartin, C., Galea, E., Lakatos, A., O’Callaghan, J. P., Petzold, G. C., Serrano-Pozo, A., Steinhäuser, C., Volterra, A., Carmignoto, G., Agarwal, A., Allen, N. J., Araque, A., Barbeito, L., Barzilai, A., Bergles, D. E., Bonvento, G., Butt, A. M., Chen, W.-T., Cohen-Salmon, M., … Verkhratsky, A. (2021). Reactive astrocyte nomenclature, definitions, and future directions. Nature Neuroscience, 24(3), 312–325. 10.1038/s41593-020-00783-4

Evans, W. R., Baskar, S. S., Ana Raquel, C. E. C., Ravoori, S., Arigbe, A., & Huda, R. (2024). Functional activation of dorsal striatum astrocytes improves movement deficits in hemi-parkinsonian mice. 10.1101/2024.04.02.587694

Fisher, T. M., & Liddelow, S. A. (2024). Emerging roles of astrocytes as immune effectors in the central nervous system. Trends in Immunology, 45(10), 824–836. 10.1016/j.it.2024.08.008

Fumagalli, M., Lombardi, M., Gressens, P., & Verderio, C. (2018). How to reprogram microglia toward beneficial functions. Glia, 66(12), 2531–2549. 10.1002/glia.23484

Garland, E. F., Hartnell, I. J., & Boche, D. (2022). Microglia and Astrocyte Function and Communication: What Do We Know in Humans? Frontiers in Neuroscience, 16. 10.3389/fnins.2022.824888

He, Y., Appel, S., & Le, W. (2001). Minocycline inhibits microglial activation and protects nigral cells after 6-hydroxydopamine injection into mouse striatum. Brain Research, 909(1–2), 187–193. 10.1016/S0006-8993(01)02681-6

Kuter, K., Kolasiewicz, W., Gołembiowska, K., Dziubina, A., Shulze, G., Berghauzen, K., Wardas, J., & Ossowska, K. (2011). Partial lesion of the dopaminergic innervation of the ventral striatum induces “depressive-like” behavior of rats. Pharmacological Reports, 63(6), 1383–1392. 10.1016/S1734-1140(11)70702-2

Kieburtz K et al. (2008). A Pilot Clinical Trial of Creatine and Minocycline in Early Parkinson Disease. Clinical Neuropharmacology, 31(3), 141–150. 10.1097/WNF.0b013e3181342f32

Kolasiewicz, W., Kuter, K., Berghauzen, K., Nowak, P., Schulze, G., & Ossowska, K. (2012). 6-OHDA injections into A8–A9 dopaminergic neurons modelling early stages of Parkinson’s disease increase the harmaline-induced tremor in rats. Brain Research, 1477, 59–73. 10.1016/j.brainres.2012.08.015

Kuter, K., Kratochwil, M., Berghauzen-Maciejewska, K., Głowacka, U., Sugawa, M. D., Ossowska, K., & Dencher, N. A. (2016). Adaptation within mitochondrial oxidative phosphorylation supercomplexes and membrane viscosity during degeneration of dopaminergic neurons in an animal model of early Parkinson’s disease. Biochimica et Biophysica Acta (BBA) - Molecular Basis of Disease, 1862(4), 741–753. 10.1016/j.bbadis.2016.01.022

Kuter, K., Olech, Ł., & Głowacka, U. (2018). Prolonged Dysfunction of Astrocytes and Activation of Microglia Accelerate Degeneration of Dopaminergic Neurons in the Rat Substantia Nigra and Block Compensation of Early Motor Dysfunction Induced by 6-OHDA. Molecular Neurobiology, 55(4), 3049–3066. 10.1007/s12035-017-0529-z

Kuter, K., Olech, Ł., Głowacka, U., & Paleczna, M. (2019). Astrocyte support is important for the compensatory potential of the nigrostriatal system neurons during early neurodegeneration. Journal of Neurochemistry, 148(1), 63–79. 10.1111/jnc.14605

Kuter, K., Śmiałowska, M., Wierońska, J., Zięba, B., Wardas, J., Pietraszek, M., Nowak, P., Biedka, I., Roczniak, W., Konieczny, J., Wolfarth, S., & Ossowska, K. (2007). Toxic influence of subchronic paraquat administration on dopaminergic neurons in rats. Brain Research, 1155, 196–207. 10.1016/j.brainres.2007.04.018

Kuter, K. Z., Olech, Ł., Głowacka, U., & Paleczna, M. (2021). Increased Beta-Hydroxybutyrate Level Is Not Sufficient for the Neuroprotective Effect of Long-Term Ketogenic Diet in an Animal Model of Early Parkinson’s Disease. Exploration of Brain and Liver Energy Metabolism Markers. International Journal of Molecular Sciences, 22(14), 7556. 10.3390/ijms22147556

Landim Falcão Tavares Ferreira, P., & De Barros Viana, G. S. (2015). Minocycline exerts a neuroprotective action against 6-OHDA-induced neurotoxicity: in vivo and in vitro studies. Neurology and Neuroscience. 10.3823/348

Li, Q., & Barres, B. A. (2018). Microglia and macrophages in brain homeostasis and disease. Nature Reviews Immunology, 18(4), 225–242. 10.1038/nri.2017.125

Markulin, I., Matasin, M., Turk, V. E., & Salković-Petrisic, M. (2022). Challenges of repurposing tetracyclines for the treatment of Alzheimer’s and Parkinson’s disease. Journal of Neural Transmission, 129(5–6), 773–804. 10.1007/s00702-021-02457-2

Nagai, J., Yu, X., Papouin, T., Cheong, E., Freeman, M. R., Monk, K. R., Hastings, M. H., Haydon, P. G., Rowitch, D., Shaham, S., & Khakh, B. S. (2021). Behaviorally consequential astrocytic regulation of neural circuits. Neuron, 109(4), 576–596. 10.1016/j.neuron.2020.12.008

Ossowska, K., Wardas, J., Śmiałowska, M., Kuter, K., Lenda, T., Wierońska, J. M., Zięba, B., Nowak, P., Dąbrowska, J., Bortel, A., Kwieciński, A., & Wolfarth, S. (2005). A slowly developing dysfunction of dopaminergic nigrostriatal neurons induced by long-term paraquat administration in rats: an animal model of preclinical stages of Parkinson’s disease? European Journal of Neuroscience, 22(6), 1294–1304. 10.1111/j.1460-9568.2005.04301.x

Pampuscenko, K., Jankeviciute, S., Morkuniene, R., Sulskis, D., Smirnovas, V., Brown, G. C., & Borutaite, V. (2025). S100A9 protein activates microglia and stimulates phagocytosis, resulting in synaptic and neuronal loss. Neurobiology of Disease, 206, 106817. 10.1016/j.nbd.2025.106817

Paulsen, R. E., Contestabile, A., Villani, L., & Fonnum, F. (1987). An In Vivo Model for Studying Function of Brain Tissue Temporarily Devoid of Glial Cell Metabolism: The Use of Fluorocitrate. Journal of Neurochemistry, 48(5), 1377–1385. 10.1111/j.1471-4159.1987.tb05674.x

Purushotham, S. S., & Buskila, Y. (2023). Astrocytic modulation of neuronal signalling. Frontiers in Network Physiology, 3. 10.3389/fnetp.2023.1205544

Quintero, E. M., Willis, L., Singleton, R., Harris, N., Huang, P., Bhat, N., & Granholm, A.-C. (2006). Behavioral and morphological effects of minocycline in the 6-hydroxydopamine rat model of Parkinson’s disease. Brain Research, 1093(1), 198–207. 10.1016/j.brainres.2006.03.104

Ravina B. (2006). A randomized, double-blind, futility clinical trial of creatine and minocycline in early Parkinson disease. Neurology, 66(5), 664–671. 10.1212/01.wnl.000020125257661.e1

Robinson, T. E., & Whishaw, I. Q. (1988). Normalization of extracellular dopamine in striatum following recovery from a partial unilateral 6-OHDA lesion of the substantia nigra: a microdialysis study in freely moving rats. Brain Research, 450(1–2), 209–224. 10.1016/0006-8993(88)91560-0

Robinson, T., Mocsary, Z., Camp, D., & Whishaw, I. (1994). Time course of recovery of extracellular dopamine following partial damage to the nigrostriatal dopamine system. The Journal of Neuroscience, 14(5), 2687–2696. 10.1523/JNEUROSCI.14-05-02687.1994

Sriram, K., Miller, D. B., & O’Callaghan, J. P. (2006). Minocycline attenuates microglial activation but fails to mitigate striatal dopaminergic neurotoxicity: role of tumor necrosis factor-α. Journal of Neurochemistry, 96(3), 706–718. 10.1111/j.1471-4159.2005.03566.x

Stefanova, N., Mitschnigg, M., Ghorayeb, I., Diguet, E., Geser, F., Tison, F., Poewe, W., & Wenning, G. K. (2004). Failure of neuronal protection by inhibition of glial activation in a rat model of striatonigral degeneration. Journal of Neuroscience Research, 78(1), 87–91. 10.1002/jnr.20233

Sun, C., Wang, Y., Mo, M., Song, C., Wang, X., Chen, S., & Liu, Y. (2019). Minocycline Protects against Rotenone-Induced Neurotoxicity Correlating with Upregulation of Nurr1 in a Parkinson’s Disease Rat Model. BioMed Research International, 2019, 1–7. 10.1155/2019/6843265

Tomás-Camardiel, M., Rite, I., Herrera, A. J., de Pablos, R. M., Cano, J., Machado, A., & Venero, J. L. (2004). Minocycline reduces the lipopolysaccharide-induced inflammatory reaction, peroxynitrite-mediated nitration of proteins, disruption of the blood–brain barrier, and damage in the nigral dopaminergic system. Neurobiology of Disease, 16(1), 190–201. 10.1016/j.nbd.2004.01.010

Tremblay, M.-E., Cookson, M. R., & Civiero, L. (2019). Glial phagocytic clearance in Parkinson’s disease. Molecular Neurodegeneration, 14(1), 16. 10.1186/s13024-019-0314-8

Ugalde-Muñiz, P., Fetter-Pruneda, I., Navarro, L., García, E., & Chavarría, A. (2020). Chronic Systemic Inflammation Exacerbates Neurotoxicity in a Parkinson’s Disease Model. Oxidative Medicine and Cellular Longevity, 2020, 1–19. 10.1155/2020/4807179

Voronkov, D., Stavrovskaya, A., Olshanskiy, A., Guschina, A., Khudoerkov, R., & Illarioshkin, S. (2021). The Influence of Striatal Astrocyte Dysfunction on Locomotor Activity in Dopamine-depleted Rats. Basic and Clinical Neuroscience Journal, 12(6), 767–776. 10.32598/bcn.2021.1923.1

Wang, Y., Wang, Q., Yu, R., Zhang, Q., Zhang, Z., Li, H., Ren, C., Yang, R., & Niu, H. (2020). Minocycline inhibition of microglial rescues nigrostriatal dopaminergic neurodegeneration caused by mutant alpha-synuclein overexpression. Aging, 12(14), 14232–14243. 10.18632/aging.103440

Wolf, S. A., Boddeke, H. W. G. M., & Kettenmann, H. (2017). Microglia in Physiology and Disease. Annual Review of Physiology, 79(1), 619–643. 10.1146/annurev-physiol-022516-034406

Wu, D. C., Jackson-Lewis, V., Vila, M., Tieu, K., Teismann, P., Vadseth, C., Choi, D.-K., Ischiropoulos, H., & Przedborski, S. (2002). Blockade of Microglial Activation Is Neuroprotective in the 1-Methyl-4-Phenyl-1,2,3,6-Tetrahydropyridine Mouse Model of Parkinson Disease. The Journal of Neuroscience, 22(5), 1763–1771. 10.1523/JNEUROSCI.22-05-01763.2002

Yang, L., Sugama, S., Chirichigno, J. W., Gregorio, J., Lorenzl, S., Shin, D. H., Browne, S. E., Shimizu, Y., Joh, T. H., Beal, M. F., & Albers, D. S. (2003). Minocycline enhances MPTP toxicity to dopaminergic neurons. Journal of Neuroscience Research, 74(2), 278–285. 10.1002/jnr.10709

